# Early developmental exposure to Fluoxetine and Citalopram results in different neurodevelopmental outcomes

**DOI:** 10.1101/780411

**Authors:** Karine Liu, Alfonso Garcia, Jenn J. Park, Alexis A. Toliver, Lizmaylin Ramos, Carlos D. Aizenman

## Abstract

Although selective serotonin reuptake inhibitors (SSRIs) are commonly prescribed for prenatal depression, there exists controversy over the adverse effects of SSRI use on fetal development. Few studies have adequately isolated outcomes due to SSRI exposure and those due to maternal psychiatric conditions. Here, we directly investigated the outcomes of exposure to widely-used SSRIs fluoxetine and citalopram on the developing nervous system of *Xenopus laevis* tadpoles, using an integrative experimental approach. We exposed tadpoles to low doses of citalopram and fluoxetine during a critical developmental period and found that the different groups of tadpoles displayed opposing behavioral effects. While both groups showed reduced schooling behavior, the fluoxetine group showed increased seizure susceptibility and reduced startle habituation. In contrast, the citalopram treated tadpoles had decreased seizure susceptibility and increased habituation. Both groups had abnormal dendritic morphology in the optic tectum, a brain area important for all three behaviors tested. Whole-cell electrophysiological recordings of tectal neurons showed no differences in synaptic function across groups; however, tectal cells from fluoxetine-treated tadpoles had decreased voltage gated K+ currents while cells in the citalopram group had increased K+ currents. Both the behavior and electrophysiological findings indicate that cells and circuits in the fluoxetine treated optic tecta are hyperexcitable, while the citalopram group exhibits decreased excitability. Taken all together, these results show that early developmental exposure to SSRIs is sufficient to induce neurodevelopmental effects, however these effects can be complex and vary depending on the SSRI used. This may explain some of the discrepancies across human studies, and further underscores the importance of serotonergic signaling for the developing nervous system.

## Introduction

The rate of utilization of antidepressant medications to treat major depressive disorder during pregnancy has continuously risen in the past decade (Cooper et al., 2007; Diav-Citrin and Ornoy, 2012; Gavin et al., 2005). One of the more commonly-prescribed class of antidepressants, selective serotonin-reuptake inhibitors (SSRIs), have a principal function of increasing serotonin concentration in the brain, and because these drugs can permeate the placenta and the developing fetal blood brain barrier (Hendrick et al., 2003; Rampono et al., 2004), there is opportunity for these pharmacological interventions to influence fetal brain development. The long-term consequences, including the risk of developing neurodevelopmental disorders in children exposed to SSRIs in utero, are not well understood, and an etiological connection between developmental levels of serotonin and autism spectrum disorders (ASD) has long been suggested.

Work done over five decades ago first pointed to hyperserotonemia in a subset of ASD children (Schain and Freedman, 1961). Today, this hypothesis has held true, with nearly 1 in 3 ASD children demonstrating 50-70% higher blood serotonin concentration (Whitaker-Azmitia, 2001, 2005). Although serotonin cannot bypass the mature blood brain barrier, it can permeate the BBB during earlier stages and especially prenatally, as such increases in maternal serotonin can affect the fetal brain. Serotonin is one of the first neurotransmitter systems to emerge in the developing brain (Shah et al., 2018) and serotonergic activity is critical for the formation of neural circuitry during perinatal development (Brummelte et al., 2017; Lauder and Bloom, 1974). While alterations in normal serotonin levels are believed to result in adverse neurodevelopmental outcomes (Shah et al., 2018).

Various studies have looked for positive correlations between increased prevalence of ASD and *in utero* SSRI exposure. In a large meta-analysis of environmental risk factors for ASD, SSRI exposure was one of the associations that retained high level of significance (Kim et al., 2019). However, recent comprehensive reviews (Millard et al., 2017; Rotem-Kohavi and Oberlander, 2017), describe the lack of consensus among these clinical studies and meta-analyses. While some report an increased risk of ASD following prenatal SSRI exposure (Boukhris et al., 2016; Croen et al., 2011; Kim et al., 2019), others have countered such conclusions (Malm et al., 2016; Sorensen et al., 2013; Suri et al., 2011). These inconsistencies in human studies, confounded by various factors such as the presence of psychiatric illness, or length of studies, highlight the need for a mechanistic approach to test whether early developmental exposure to SSRIs alone is sufficient to induce neurodevelopmental effects.

In rodents, prenatal and perinatal exposure to fluoxetine (Ansorge et al., 2004; Gemmel et al., 2017; Olivier et al., 2011; Sprowles et al., 2017), and citalopram (Simpson et al., 2011; Sprowles et al., 2017) resulted in pups with decreased exploratory behavior, increased latency to begin feeding, enhanced anxiety, repetitive behaviors, altered HPA axis function and decreased social play. Furthermore, plasma fluoxetine transfer from mother to pup was around 83%. Despite these behavioral studies, there are almost no studies looking directly at the effects on neural circuit function of early SSRI exposure (Yu et al., 2019).

Here we take an integrative approach which uses a series of electrophysiological, neuroanatomical and behavioral approaches in *Xenopus laevis* tadpoles to directly study the neurodevelopmental effects of early exposure to SSRIs. Xenopus tadpoles have been used to successfully model neurodevelopmental disorders (Pratt and Khakhalin, 2013) with early exposure to valproic acid (VPA), an anti-epileptic drug associated with higher incidence of ASD following pre-natal exposure, inducing neurodevelopmental deficits (James et al., 2015). Furthermore, serotonin is an important regulator of CNS development in amphibians (De Lucchini et al., 2003; Sillar et al., 2002), suggesting that Xenopus tadpoles are a good model to test the effects of developmental SSRI exposure on circuit function. Using these assays, we tested whether early exposure to Fluoxetine and Citalopram results in functional abnormalities associated with neurodevelopmental disorders, and found that they both affect neural circuit development but in distinct ways.

## Results

In this study we compare three treatment groups: naive controls, tadpoles reared between developmental stages 42/43 through 49 in 1.25 mg/L Fluoxetine and in 2.5 mg/L Citalopram. Concentrations were determined based on a survival curve, and were chosen such that we used the highest concentration at which all animals survived for 10 days without any adverse health effects. To test whether exposure to Fluoxetine or Citalopram induces neurodevelopmental deficits, we performed a series of behavioral tests that are known to be altered in our VPA-treated tadpoles, including schooling, seizure susceptibility and startle habituation.

We had previously described a deficit in schooling behavior in tadpoles reared in VPA (James et al., 2015). When placed in a large arena, tadpoles exhibit schooling behavior that depends on visual, mechanosensory and olfactory cues (Katz et al., 1981; Lum et al., 1982). Thus, impairment of schooling may indicate impaired sensorimotor integration and/or impaired social behavior. Tadpoles were placed in groups of 15-17 in an arena and allowed to organize themselves into clusters. Control tadpoles swim in groups and co-orient with each other. To distinguish any abnormalities in schooling, inter-tadpole angles and inter-tadpole distances from treated tadpoles were compared to controls. While both SSRIs resulted in differences in schooling, we found qualitative differences between the two different SSRIs. Tadpoles reared in Citalopram showed a significant shift toward larger inter tadpole distances and a more random distribution of inter tadpole angles (Fig. 1A,B left panels, 1C, KS test for distances: p=0.0004, and angles: p<0.0001, n=12 experiments for each group), consistent with decreased schooling and similar to what was observed in VPA treated tadpoles (James et al., 2015). In contrast, Fluoxetine-treated tadpoles exhibited a significant shift towards smaller inter tadpole distances as well as a random distribution of inter tadpole angles (Fig. 1A,B right panels, 1C, KS test for distances: p<0.0001, and angles: p<0.0001, n=17 experiments for each group). The reason for the tighter clusters of tadpoles was not because of enhanced schooling as these clusters were not organized groups of tadpoles swimming in the same direction, but rather due to the fact that tadpoles appeared to swim randomly and tended to stop swimming once they collided with another tadpole, forming static groups of tadpoles (Fig. 1C). Thus, while exposure to both SSRIs resulted in abnormal schooling behavior, these differences have distinct profiles, suggesting that Citalopram and Fluoxetine may have different mechanisms of action.

**Figure 1.**
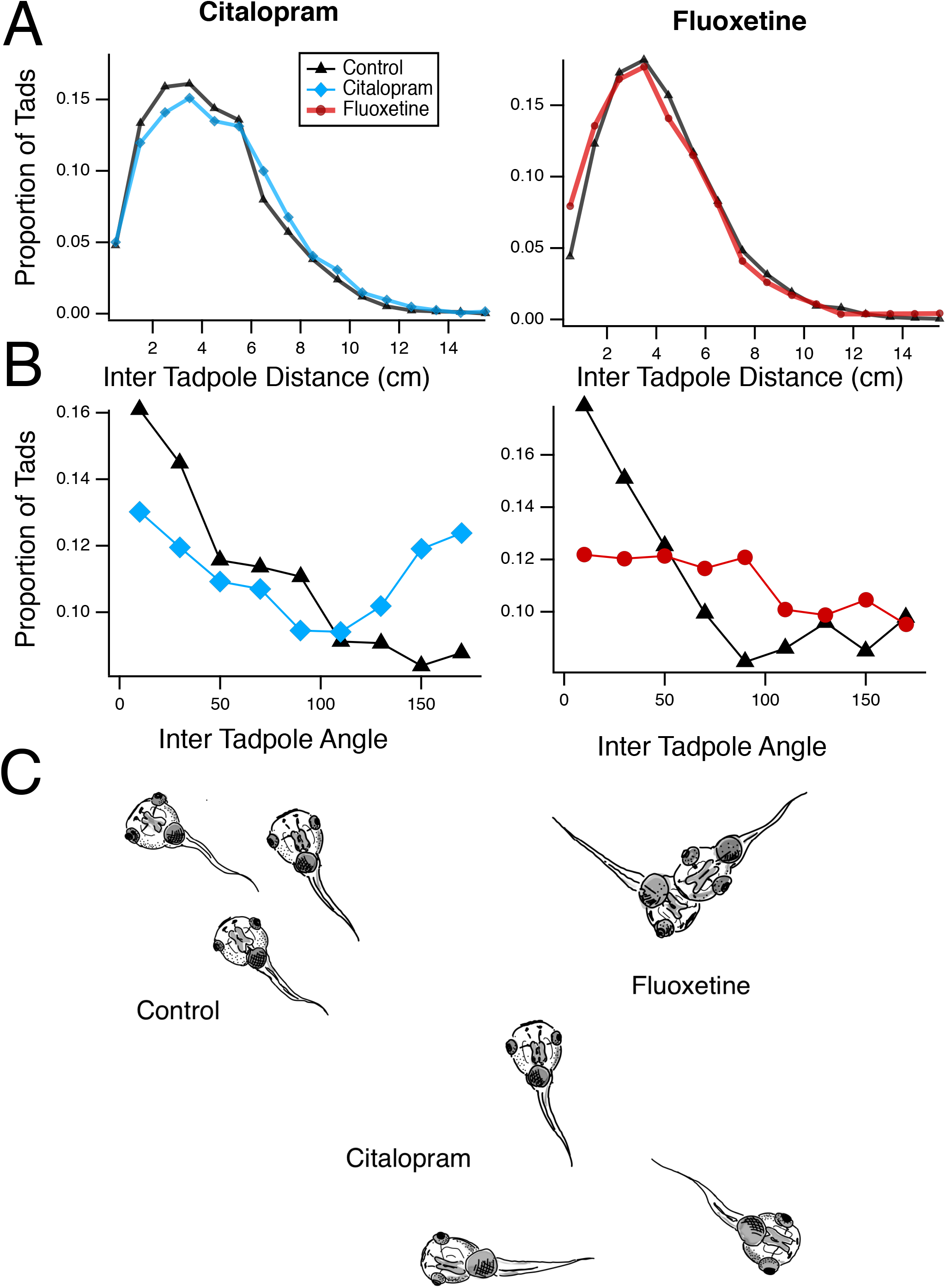
Early SSRI treatment results in disrupted schooling behavior. **A.** Distribution of inter-tadpole distances for the Citalopram (left) and Fluoxetine (right) – treated groups, compared to controls. The Citalopram group shows a significant shift toward longer distances, indicating more disperse swimming patterns and decreased schooling. The fluoxetine group shows a shift toward smaller distances, indicating increased clustering of animals. **B.** Distribution of inter-tadpole angles. Control groups show a larger number of tadpoles with similar angles, indicating that they are swimming in the same direction. Both experimental groups show a broader distribution of inter-tadpole angles, indicating that tadpoles are swimming in multiple directions relative to each other and thus do not have organized schooling. **C.** Schematic illustration of the different effects of Fluoxetine and Citalopram treatment on school organization.

Next, we tested seizure susceptibility. When exposed to a convulsant agent (pentenyltetrazole, PTZ at 5 mM) tadpoles will develop periodic seizures within a few minutes (Bell et al., 2011). VPA-treated tadpoles are known to exhibit hyper-connectivity within their CNS which results in a higher susceptibility to PTZ-induced seizures, as measured by an increase in the frequency of seizure episodes (James et al., 2015). We compared seizure frequency across treatment groups and found that Fluoxetine treated tadpoles had an overall higher frequency of seizure episodes compared to controls (Fig. 2A, Flx 1.04±0.04 seizures/min, n=57, Ctrl 0.78±0.04 seizures/min, n=80), while Citalopram treated tadpoles had a lower seizure frequency (0.7±0.05 seizures/min, n=35). This difference became increasingly larger and significant over the 20 minute exposure window (p<0.0001 for Treatment effect, 2-way ANOVA), in which the seizure frequency in the Fluoxetine group increased noticeably, with little or no change in the Citalopram group over time (Fig. 2B). These differences became significantly different between treatment groups and controls by 20 minutes. Another measure that reflects neuronal hyper-connectivity is startle-habituation. Startle habituation is measured by placing tadpoles in an arena and exposing them to auditory clicks which trigger a startle response. Tadpoles will show rapid habituation of the startle response when presented with a series of sound clicks presented every 5 seconds, and when this series is repeated 5 minutes later, they retain this habituation, referred to as short-term habituation. Even after a final rest interval of 15 minutes, the tadpoles remain habituated, this measure is referred to as long term habituation. Click intensity was calibrated so that the control group showed a small amount of short and long-term habituation, allowing us to better measure changes in either direction. The Fluoxetine treated group showed significantly enhanced startle responses, compared to controls, that persisted even across various stimulus bouts (Fig. 2C, p<0.0001 2-way ANOVA with Dunnet’s multiple comparison test, n= 24 for each group). In contrast Citalopram treated tadpoles showed a much stronger short and long-term habituation profile across bouts (p=<0.0001 2-way ANOVA with Dunnet’s multiple comparison test).

**Figure 2.**
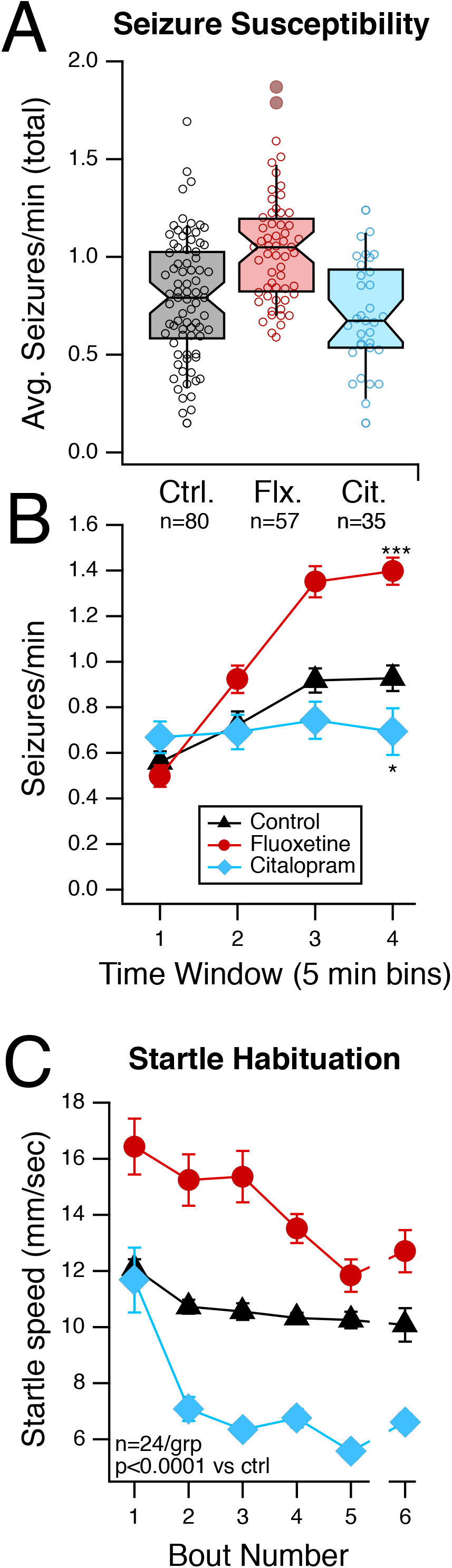
Early SSRI treatment alters seizure susceptibility and startle habituation. **A.** Distribution of number of seizures per minute during the entire 20 minute observation period, for control and experimental groups. Individual circles represent one tadpole, box represents median and IQR, whiskers represent 9^th^ and 91^st^ percentiles. The notch represents the 95% confidence interval. **B.** Plot shows the average (±SEM) number of seizures/minute for each 5 minute interval during the observation period. Fluoxetine shows a gradual significant increase in seizure frequency while citalopram shows a decrease. *p< 0.05, ***p<0.001 **C.** Average startle speeds for each stimulus bout shows that the Fluoxetine treated group shows decreased habituation and overall more responsiveness to startles over time, while the Citalopram group shows strong habituation.

Both seizure and startle data indicate that Fluoxetine-treated tadpoles show behaviors consistent with hyper-connectivity and increased excitability, while Citalopram-treated tadpoles show pronounced habituation and decreased seizure susceptibility. These differences may stem from different effects of SSRI exposure on neural circuit development. All three behaviors tested involve activation of tectal circuitry (Bell et al., 2011; James et al., 2015), and thus we performed a series of neuroanatomical and electrophysiological experiments in the tectum in all three treatment groups. To see if differences in tectal circuit function could account for the behavioral effects of early SSRI exposure. First, we examined whether the SSRI-exposed groups showed any abnormalities in tectal neuron dendritic architecture. We used sparse electroporation of GFP-expressing plasmids (see methods; (Dhande et al., 2011)) to drive expression of GFP in individual tectal neurons, and after the SSRI exposure period we used *in vivo* confocal microscopy to image these cells. We then reconstructed the morphology of individual neurons and performed a 3D Scholl analysis to measure dendritic branch distribution. Both Fluoxetine and Citalopram groups showed a significant difference in overall branching patterns (Fig. 3A), such that the dendritic arbors showed fewer branch points all along the length of the primary dendrites. Both citalopram and fluoxetine groups showed a significantly different distribution of branch points from controls (Fig. 3B, p<0.001 KS Test), although there was no difference in the total dendritic branch length (Average TDBL, Control: 667±139 µm, n=15, Citalopram: 611±132 µm, n=16, Fluoxetine: 628±113 µm, n=9). This indicates that while both SSRIs are affecting underlying circuits, their effect on dendritic architecture is not sufficient to explain their differential effects on behavior.

**Figure 3.**
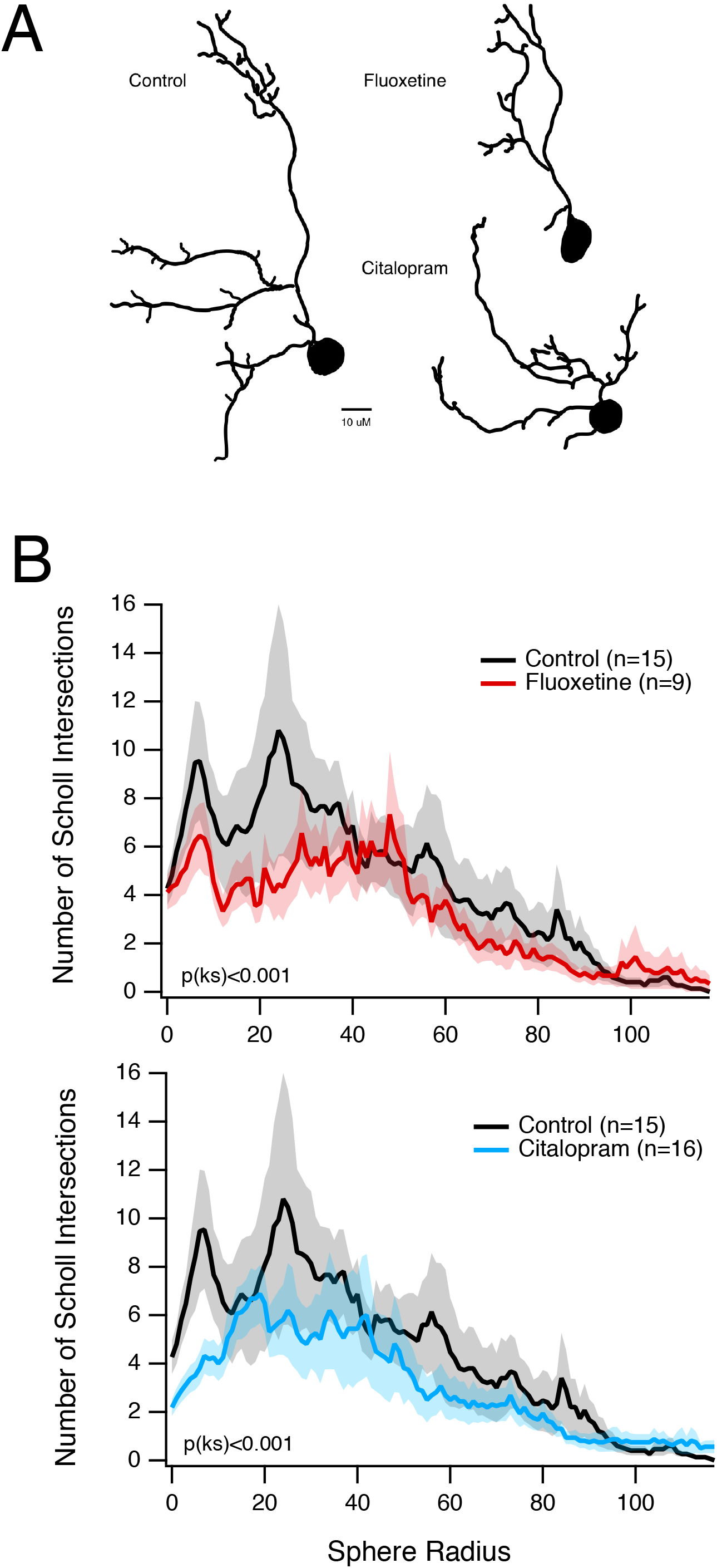
Early SSRI exposure alters normal development of tectal neuron dendritic morphology. **A.** Sample 2D tracings of 3D neuronal reconstructions from all three experimental groups showing overall smaller dendritic fields for both treated groups. **B.** Averaged 3D Scholl analyses showing branch distribution as a function of distance from the soma. Lighter colored areas indicate SEM at each location. SSRI-treated groups show an impoverished branching pattern, compared to controls.

We next performed a series of whole-cell voltage clamp experiments to measure synaptic properties in optic tectal neurons using an *ex vivo* whole-brain preparation (Wu et al., 1996). One possibility is that early SSRI exposure caused hyperconnectivity as seen in VPA treated tadpoles (James et al., 2015). These would be reflected as changes in the frequency of spontaneous excitatory postsynaptic currents (sEPSC). We found that across treatment groups, SSRI exposure had no significant effects on sEPSC amplitudes or frequency (Fig. 4A, Control, amplitude: 6±0.2 pA, frequency: 4.2±0.4 events/sec, n=48; Fluoxetine, amplitude: 6.5±0.4 pA, frequency: 4.6±0.5 events/sec, n=37; Citalopram, amplitude: 8.1±0.6 pA, frequency: 5.9±0.8 events/sec, n=36). Another possibility is that SSRI exposure could result in an imbalance of excitation to inhibition, as described in some autism models (Lee et al., 2017). We used a stimulating electrode placed on the optic nerve to evoke afferent excitatory inputs and feedforward inhibition. These were isolated by holding the membrane potential at the GABA (−45 mV) and Glutamate (+5 mV) receptor reversal potentials and then measuring the peak responses (Fig. 4B). We found no difference in the Excitation/Inhibition ratio across groups (Control, E/I ratio: 0.68±0.1, n=23; Fluoxetine: 0.67±0.08, n=30; Citalopram: 0.5±0.05, n=16). Likewise, we also found no difference in paired-pulse facilitation, which could indicate a change in presynaptic release probability of the visual inputs (Control, PPF ratio: 1.4±0.2, n=18; Fluoxetine: 1.3±0.2, n=18; Citalopram: 1.5±0.1, n=17).

**Figure 4.**
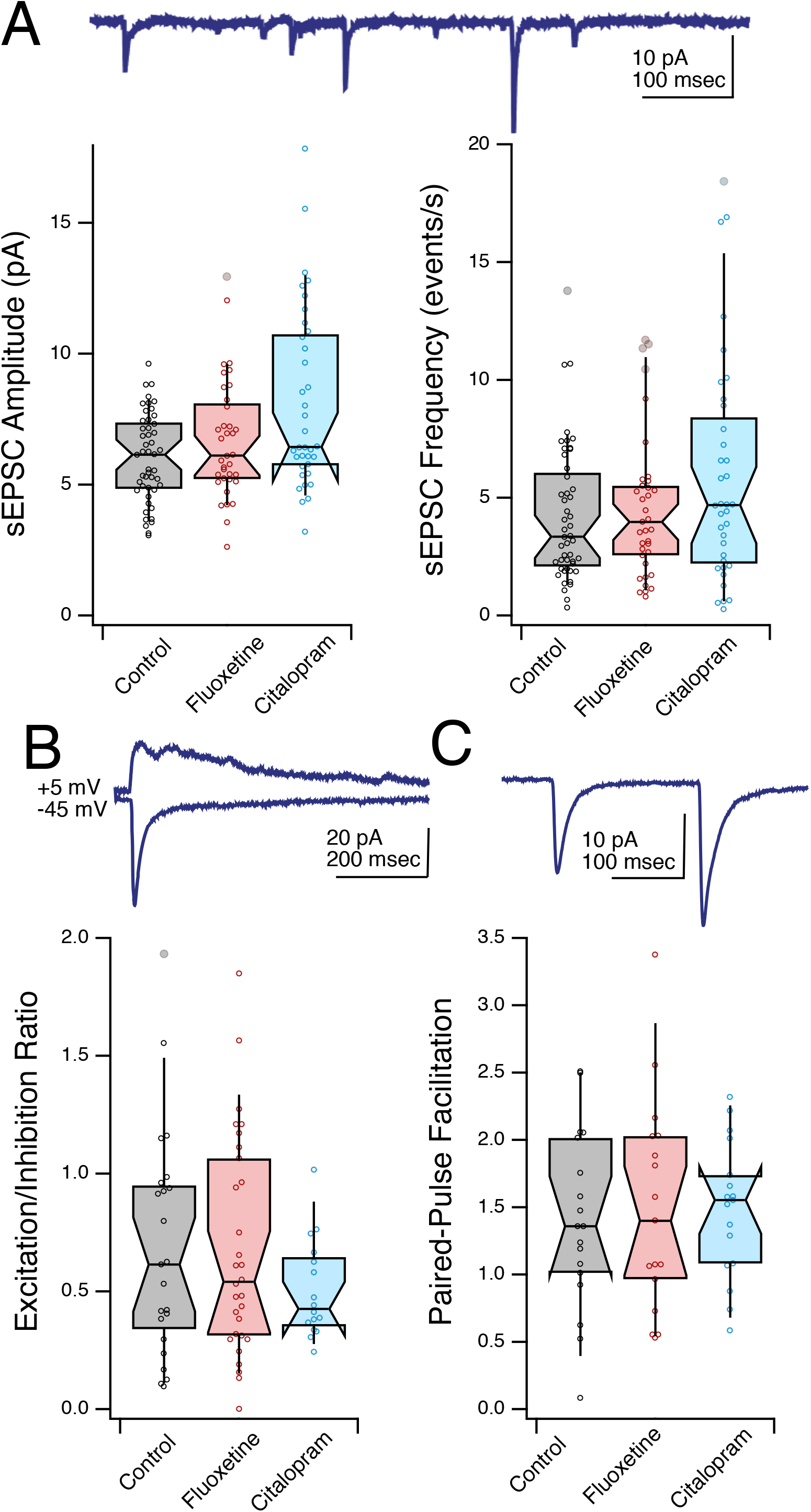
SSRI exposure during early development does not alter synaptic transmission in the tectum. **A.** Box plot shows spontaneous EPSC amplitude (left) and frequency (right) for the different experimental groups. Individual circles represent one cell, box represents median and IQR, whiskers represent 9^th^ and 91^st^ percentiles. The notch represents the 95% confidence interval. Inset shows sample recording of sEPSCs. **B.** Box plot shows the excitation and inhibition ratio as measured by dividing the peak excitatory response recorded at −45 mV and the peak inhibitory response recorded at +5 mV (inset). **C.** Box plot shows the amount of paired pulse facilitation as calculated by the ratio of the second response over the first response (inset).

To measure intrinsic excitability, we performed a series current-voltage experiments to measure voltage-gated Na+ and K+ currents (Fig. 5A,B, (Ciarleglio et al., 2015; Pratt and Aizenman, 2007). While we found no differences in peak voltage-gated Na+ currents (Fig. 5C, K+ current Control: 338±29 pA, n=52; Fluoxetine:327±30 pA, n=51; Citalopram: 337±17 pA, n=39) there were significant differences in peak voltage-gated K+ currents, such that the Fluoxetine group had decreased K+ currents (Fig. 5D, K+ current Control: 1950±146 pA, n=52; Fluoxetine:1329±136 pA, n=51; p<0.01 Dunnet), while the Citalopram group exhibited larger K+ currents (Citalopram: 2656±105 pA, n=39, p<0.001 Dunnet). Decreased K+ currents would mean that cells in the Fluoxetine group would have greater intrinsic excitability, while the Citalopram treated cell would have lower excitability, consistent with the behavioral findings where the Fluoxetine treated tadpoles had more seizures and an exaggerated startle response.

**Figure 5.**
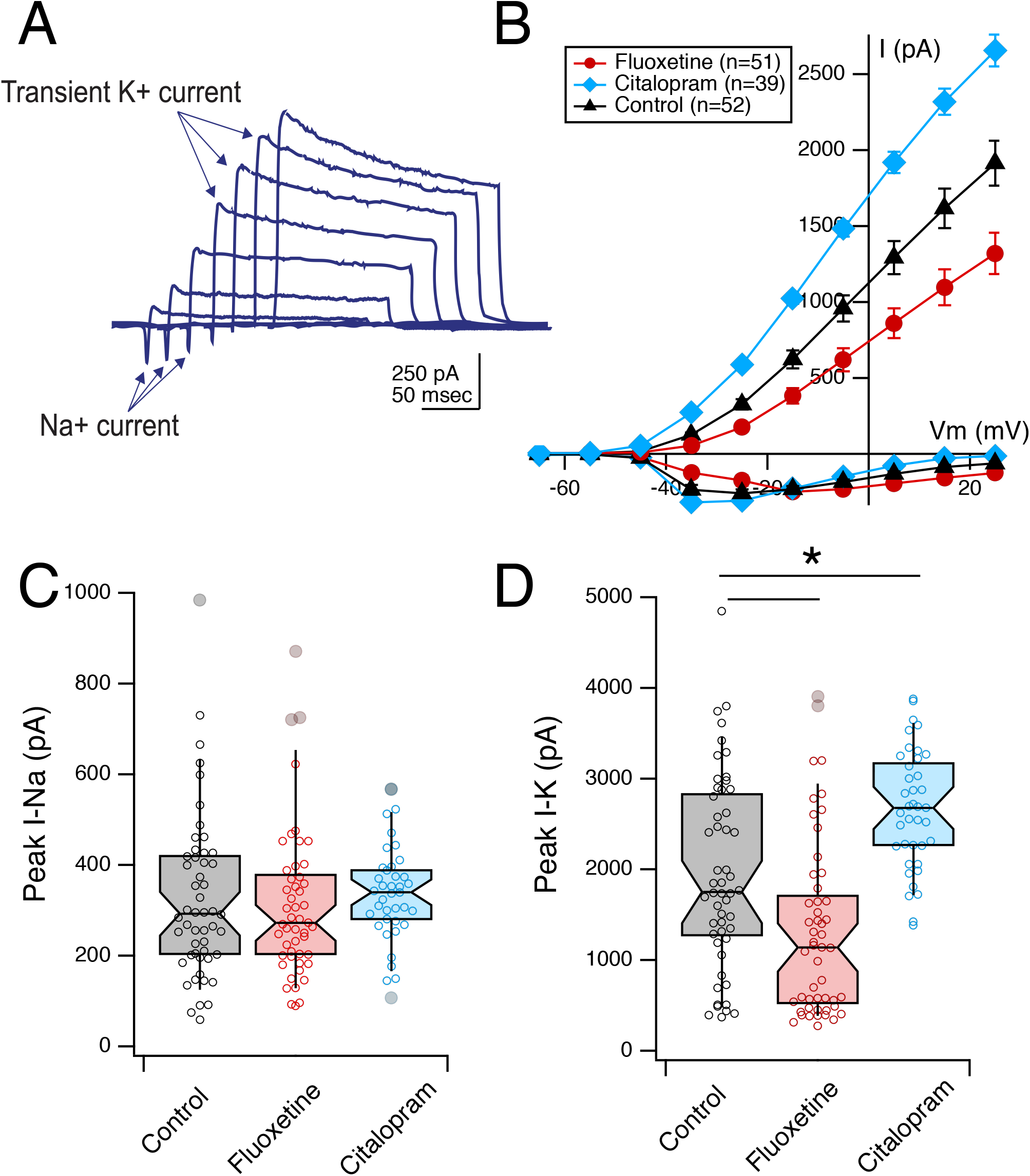
Early exposure to Fluoxetine and Citalopram results in opposite effects on voltage-gated K+ currents. **A.** Representative traces showing a family of current recordings in response to a series of depolarizing steps. **B.** Averaged current-voltage plots for the Na+ and K+ currents across experimental groups. Notice the enhanced outward K+ current in the Citalopram cells and the decreased current in the Fluoxetine cells. Error bars are SEM. **C, D.** Box plots denting the peaks Na+ and K+ currents across groups. *p<0.05. Individual circles represent one cell, box represents median and IQR, whiskers represent 9^th^ and 91^st^ percentiles. The notch represents the 95% confidence interval.

## Discussion

Our data here investigate the impact of early exposure to SSRIs in the developing Xenopus laevis optic tectum. We observed opposing behavioral outcomes between fluoxetine-treated and citalopram-treated animals. While both SSRI-exposed groups failed to show normal schooling behavior the patterns by which schooling was disrupted was different. Moreover, fluoxetine-treated animals demonstrated a reduced capacity to habituate to an acoustic stimulus, but citalopram-treated animals habituated to the same high-intensity stimulus faster. In an assay of seizure frequency, fluoxetine-treated animals yielded a significant increase in seizure frequency as compared to control animals, while the citalopram groups showed a decreased seizure frequency. Both treatment groups showed overall decreased dendritic branching patterns in optic tectal neurons indicating alterations in neural connectivity. Electrophysiologically, we found no significant differences in spontaneous synaptic transmission or presynaptic facilitation. However, we did find decreased K+ currents in fluoxetine-treated neurons, consistent with increased cellular excitability, while increases in the K+ currents in citalopram-treated neurons were consistent with decreased cellular excitability. Taken together these data show that early exposure to SSRIs is sufficient to alter the neural network excitability and architecture of the optic tectum, which in turn correlates with altered behavioral outcomes. These behavioral outcomes are consistent with our prior observations in other Xenopus models of ASD (James et al., 2015; Lee et al., 2010; Pratt and Khakhalin, 2013; Truszkowski et al., 2016).

Although fluoxetine and citalopram fall within the same class of drugs, their effects lead to opposing alterations in network excitability in the developing CNS. To interpret the difference in outcomes, we can look to the pharmacodynamics of each SSRI. Although the exact mechanisms through which fluoxetine and citalopram in mediating their action at the SERT are not well understood, variation in selectivity between fluoxetine and citalopram has been previously established (Owens et al., 2002). Fluoxetine has notable binding to other monoamine transporters, including the norepinephrine transporter and dopamine transporter (Leonard, 1995). Inhibition of the uptake of these neurotransmitters has the potential to limit what consequences we can accurately assign to the excess of serotonin, as elicited by SSRIs. In fact, little is known about the effects of excess norepinephrine and dopamine during early development. Furthermore, some SSRIs can directly bind to serotonin receptors. For example, Fluoxetine, but not other SSRIs tested, caused an increase norepinephrine and dopamine in the rat prefrontal cortex acutely (Bymaster et al., 2002). They suggest that fluoxetine binding to the 5-HT2C receptor, a receptor irrelevant to direct action by SSRIs, may be responsible for serotonin-related regulation of norepinephrine and dopamine release in the PFC. These differences among the different neuromodulators may lead to complex cellular interactions that ultimately lead to different outcomes in cell physiology.

Early alterations in serotonin are known to have effects in circuit development. The finding that both treatment groups showed altered dendritic morphology is interesting. For example, serotonin is known to inhibit growth of primary dendrites via activation of 5-HT1A receptors, but promotes growth of secondary dendrites in the hippocampus via activation of 5-HT1A and 5-HT7 (Rojas et al., 2017). And other studies show that too much serotonin results in abnormal development of visual projections into the CNS (Salichon et al., 2001; Upton et al., 1999). A more careful examination of both serotonin levels and receptor activity after chronic SSRI exposure will better help understand how the dendritic architecture is affected over the course of development when serotonin levels are elevated. The same could be said for serotonin’s other effects on neural circuit development – including neurogenesis, dendritic refinement, maintenance, and synaptic remodeling (Whitaker-Azmitia, 2001, 2005).

This study extends our knowledge into the effects of serotonin modulation to the early brain, but inspires many more questions. The outcomes detected here may be better explained through further investigation of the many serotonergic receptors, specifically those important for developmental outcomes or are modulated by repeated administration of fluoxetine and citalopram. Early exposure to selective norepinephrine reuptake inhibitors (SNRIs) and dopamine reuptake inhibitors (DRIs) in a similar protocol as in this study could provide clarity on the differing effects of fluoxetine and citalopram shown here, as would also prenatal exposure to other commonly used SSRIs. Furthermore, the behavioral measures described here all are known to involve tectal processing (Bell et al., 2011; James et al., 2015; Khakhalin et al., 2014; Lee et al., 2010), and are correlated with changes in tectal cell physiology and morphology. However, it is likely that similar changes are occurring throughout the CNS. Thus, this study uses the tectum as a readout, for more generalized changes in brain development, rather than implying that these are tectal specific changes.

In conclusion, this work substantiates the varying action between two commonly prescribed SSRIs. Early fluoxetine and early citalopram exposure both appear to influence developing neural circuits, but to opposite ends, as examined at the cellular, synaptic, and network levels. These changes to intrinsic excitability accompany differing behavioral effects resultant from the same exposure. Taken together, differing off-target mechanisms between the two SSRIs are highly implicated and should inspire further effort to better inform treatment strategies for the benefit of maternal and fetal health. These studies also indicate that chronic alterations in neuromodulators can have profound effects on early neural circuit organization that may lead to neurodevelopmental disorders later in life. The differences between neuromodulators may also help explain some of the inconsistencies observed in the clinical data that tries to establish a correlation between SSRI exposure and neurodevelopmental disorders.

## Methods

### Rearing Xenopus laevis tadpoles

Wildtype *Xenopus laevis* were bred in the animal care facility of Brown University. Following fertilization, embryos were collected and maintained in a 18ºC incubator on a 12 hr:12 hr light/dark cycle. Embryos were kept in 10% Steinberg’s solution (5.8 mM NaCl, 0.067 mM KCl, 0.034 Ca(NO_3_)_2_ • 4H_2_O, 0.083 mM MgSO_4_ • 7H_2_O, 5 mM HEPES, pH 7.4). Tadpole development was measured as delineated by Nieuwkoop and Faber (1956). At stage 42 (roughly 5 dpf), tadpoles were exposed to the following drug solutions: 1.25 mg/mL Fluoxetine-HCl, 2.5 mg/mL Citalopram-HBr (Sigma-Aldrich; St. Louis, MO) or left in control media. Dosages were determined by creating dose-response curves, and choosing the highest dose that does not cause mortality nor morphological defects (data not shown). The drug plus Steinberg’s solution were replaced every 3 days to maintain consistent drug levels and maintain solution cleanliness for tadpole health, for a total exposure of 8-10 days, when tadpoles reach Stage 49.

### Behavior

#### Acoustic Startle Habituation

Before any of the behavioral experiments, tadpoles were transferred to control media and left to sit for 1-2 hours, to recover from the possible acute action of the drugs. For acoustic startle habituation, stage 49 tadpoles were individually placed into wells of a six-well culture plate as described in **James et al. 2014**. The plate was fixed between two audio speakers (SPA2210/27; Philips) which in turn were mechanically attached to the plates to transfer vibration. Each of the wells was filled with 6.5 mL of Steinberg’s solution. To account for sound variability in sound transmission between the wells, treatments and control tadpoles were cycled throughout the wells for every trial. The acoustic stimuli consisted of a period of a 100 Hz sine wave (5ms long) which was delivered every 5 seconds. This stimulus evoked a startle response from the animals. The response is result of the acoustic and vibrational stimuli from the speakers. Each experiment consisted of 6 bouts with a 5-minute rest in between bouts 1-5. A 15-minute rest period was implemented between bouts 5 and 6. Tadpole behavior was recorded with a video camera overhead the plate and tracked with EthoVision (Noldus Information Technology). A custom MATLAB script then analyzed the coordinates generated by EthoVision. By measuring the peak speed of each tadpole 2 seconds after delivery of the stimulus, the script generated the startle velocities of the tadpoles in response to each acoustic startle.

#### Seizure Susceptibility

Seizure susceptibility and intensity serve as reliable markers of neurodevelopmental disorders that may indicate hyper connectivity in the nervous system. Treatment and control tadpoles were transferred into individual wells in a six-well plate, each of which filled with 10 mL of 5mM pentylenetrerazole (PTZ; P6500 from Sigma Aldrich) solution in 10% Steinberg’s media. The plate was diffusely illuminated from below and recorded overhead using a color digital video camera (SCB 2001; Samsung). Tadpole behavior in the wells was recorded and individually tracked by EthoVision software. Recordings lasted 20 minutes. Tracking data was then processed by an offline MATLAB script that defined the seizure events. Seizure events were defined by periods of rapid irregular movement, interrupted by certain periods of immobility. Seizure events were also defined as swimming activity greater than half of the maximal velocity of the tadpole within in each trial. We calculated each tadpole’s average seizure frequency, time to first seizure, and average seizure length. To track changes in seizure frequency over time, each trial was further divided into four 5-minute windows.

#### Schooling Behavior

To measure schooling behavior, batches of 15-17 Stage 49 tadpoles were placed in a 17 cm glass bowl, filled with 350 mL of rearing media. To ensure consistency in the data, control tadpoles matched treated-raised tadpoles in number. For each trial, a still image of tadpole distribution in the glass bowl was taken by a webcam controlled by Yawcam software every 5 minutes. Each trial lasted an hour and generated thirteen images. A translucent enclosure around the glass dish isolated the tadpoles from possible external stimuli. After 2.5 minutes following each still photo, a strong mechanic startle was delivered. The stimulus effectively startled the tadpoles to force them to redistribute (Katz et al, 1981, **James 2015**). Consecutive startles were delivered every 5 minutes. Using NIH ImageJ, coordinates of the tadpole heads and gut were tracked manually and then exported into MATLAB for further processing. The program defined inter-tadpole distance through point set triangulation and calculated the distance between tadpoles root ((x^2^ – x^1^)^2^ + (y^2^ – y^1^)^2^). Inter-tadpole angles were measured by fitting a line through the tapdole’s body and calculating the angle relative to its neighbors.

### Whole Brain Electroporation and Reconstruction of Neuronal Morphology

To label single tectal cells, stage 45-47 tadpoles were anesthetized and whole-brain electroporation was used to transfect tectal neurons with a fluorescent protein (Haas et al., 2002). Tadpoles were anesthetized with 0.02% MS-222 and, plasmid DNA was pressure injected into the brain’s middle ventricle using a glass micropipette. Two platinum electrodes (1-2 mm) were placed on the skin overlying both sides of the midbrain. The platinum electrodes were used to send current to incorporate the DNA into the tectal cells. Three electrical pulses at 50 V with an exponential decay of 70 ms were delivered to the tadpole brain. For this electroporation, pCALNL-GFP plus pCAG-Cre plasmids were incorporated. Both plasmids need to be incorporated into a tectal cell in order to get expression, yielding sparse transfection and thus allowing us to isolate individual cells for imaging (Dhande et al., 2011). After electroporation, tadpoles were immediately returned to their respective rearing media. Electroporated tadpoles were then allowed to develop until stage 49. A Zeiss 800 confocal was used to image GFP-expressing neurons. Acquired confocal images were then used to reconstruct the three-dimensional morphology of tectal cells using Imaris (Oxford Instruments). The 3D reconstructed dendritic arbors were then further analyzed to extract the following parameters: average dendrite length, average dendrite area, average dendrite branching angle, and average dendrite mean diameter and to generate Scholl analyses.

### Whole-Cell Electrophysiology

*T*adpole brains were prepared for *ex vivo* recordings upon reaching stage 49. We anesthetized the animals by submersion in 0.01% MS-222 for several minutes. Tadpoles were dissected in external solution, a HEPES-buffered saline solution (115 mM NaCl, 2-4 mM KCl, 3 mM CaCl_2_, 0.5 MgCl_2_, 5 mM HEPES, 10 mM glucose, and 0.1 mM picrotoxin; pH 7.2, 255 mOsM). The isolated brain was prepared as previously described (Wu et al., 1996; Aizenman et al., 2013). Revealing the brain required slicing through the dorsal surface of the tadpole, cutting through the pigmented skin. Membranous tissue and commissural nerve fibers are also cut in this process. Once the brain was excised from the animal, the brain was then mounted and pinned onto a silicone elastomer (Sylgard, Dow Corning, Midland, MI) block. To expose the tectal neurons, broken micropipettes were used removed the ventricular membrane with gentle suction. Cells were visualized using a Nikon E600 FN light microscope with a 60x fluorescent water-immersion objective. To evoke synaptic responses, we placed bipolar stimulating electrodes (FHC, No. CE2C7S) in the optic chiasm.

Data was collected in whole cell voltage-clamp mode. Glass micropipettes were filled with potassium gluconate-based intracellular saline (100 mM potassium gluconate, 8 mM KCl, 5 mM NaCl, 1.5 MgCl_2_, 20 mM HEPEs, 10 mM EGTA, 2 mM ATP, 0.3 mM GTP; pH 7.2, 255 mOsm). Tectal neuron recordings were limited to the middle third of the tectum so as to minimize variations in tectal neuron maturity. Signals were recorded using a Multiclamp 700B amplifier (Axon Instruments), digitized at 10 kHz using a Digidata 1322A A-D board, and acquired using the P-Clamp Clampex 10 software. AxographX software was used to analyze recordings.

